# Global genomic epidemiology, resistome, virulome and plasmidome of the Extraintestinal Pathogenic *Escherichia coli* (ExPEC) ST38 lineage

**DOI:** 10.1101/2022.05.04.490652

**Authors:** Érica L. Fonseca, Sérgio M. Morgado, Ana Carolina P. Vicente

## Abstract

Most of the extraintestinal human infections worldwide are determined by some specific Extraintestinal Pathogenic *Escherichia coli* (ExPEC) lineages, which also present a zoonotic character. One of these lineages belongs to ST38, which is considered a uropathogenic/enteroaggregative *E. coli* hybrid and a high-risk globally disseminated ExPEC. To get insights on the aspects of ST38 epidemiology and evolution as a multidrug resistant and pathogenic lineage, this study performed a global comparative genomic analysis on ST38 genomes. A total of 376 genomes recovered from environments, humans, livestock, wild and domestic animals in all continents throughout three decades were analyzed considering the ST38 phylogenomics, resistome, plasmidome, and virulome. In general, our analyses revealed that, independently of host sources and geographic origin, these genomes were distributed in two clusters comprising clonal and non-clonal genomes. Moreover, the ST38 accessory genome was not strictly associated with clusters and sub-clusters. Interestingly, the High Pathogenicity Island (HPI) prevailed in the major cluster (cluster 2), which comprised most of the genomes from human origin recovered worldwide (2000 to 2020). In addition, the ExPEC ST38 harbors a huge and diverse plasmidome that might contribute to the evolution of this lineage, reflecting in ST38 ability to live in a diversity of niches as a commensal or pathogenic organism.

## 1. Introduction

*Escherichia coli* is a commensal bacterium and a versatile pathogen capable of causing intestinal and extraintestinal infections (Vila et al., 2016). Extraintestinal pathogenic *E. coli* (ExPEC) determines most of the human extraintestinal infections (Manges et al., 2019), and includes uropathogenic *E. coli* (UPEC), neonatal meningitis *E. coli* (NMEC), sepsis-associated *E. coli* (SEPEC), and avian pathogenic *E. coli* (APEC) (Kaper et al., 2004). ExPEC comprises many lineages that affect not only humans but also animal health, besides occurring in the environment. However, a subset of lineages/sequence types (STs) is responsible for the majority of extraintestinal pathologies in humans and animals. The recent emergence of multidrug-resistant ExPEC lineages (ST131, ST38, ST405, and ST648) raised concerns on how *E. coli* evolves and diversifies in terms of virulence and antibiotic resistance (Dale and Woodford, 2015).

The ST38 is considered a uropathogenic/enteroaggregative *E. coli* hybrid (Chattaway et al., 2004), which also presents a zoonotic character since it is often identified in natural environments and animals (Hayashi et al., 2018; de Carvalho et al., 2020; Fernandes et al., 2020; Lopes et al., 2021; Salgado-Caxito et al., 2021).

The Extend Spectrum β-Lactamase (ESBL)-coding *bla*_CTX-M_ gene is prevalent among ExPEC lineages, such as ST131, which has been associated with the global dissemination of this gene (Bevan et al., 2017; Shaik et al., 2017; Mostafa et al., 2020). Although *bla*_CTX-M_ genes have been found in ST38 (Peirano et al., 2012; Pitout, 2012; Shaik et al., 2017), this lineage is far less characterized than ST131 (Cameron et al., 2021). Therefore, to get insights on ST38 and *bla*_CTX-M_ dissemination, as well as, other features involved with its high-risk character in clinics, a global genomic analysis of ST38 should be performed taking into account the three axes of the One Health Concept (humans, animals, and natural environments), and the virulence, adaptability, and persistence traits.

Thus, this study performed a large-scale comparative genomic analysis on ST38 genomes from environments, humans, livestock, wild and domestic animals recovered from all continents throughout three decades, and determined the global ST38 phylogenomics, resistome, plasmidome, and virulome. In general, our analyses corroborated the zoonotic character of the ExPEC ST38 since it was identified clonal genomes from both animal and human sources recovered worldwide. Interestingly, it was verified that a virulence trait present in most of the ST38 genomes, the High Pathogenicity Island (HPI) from *Yersinia pestis*, prevailed in ST38 from human origin. In addition, the ExPEC ST38 was characterized by a huge and diverse plasmidome that might contribute to the evolution of this lineage, reflecting in ST38 ability to in a diversity of niches as a commensal or pathogenic organism.

## 2. Materials and Methods

### 2.1. EnteroBase ST38 meta-analysis

EnteroBase is integrated software that supports the identification of global population structures within several bacterial genera (Zhou et al., 2020). Here, a metaanalysis based on the geographical information of ST38 entries included in EnteroBase (https://enterobase.warwick.ac.uk) was performed (last updated in September 2021). The ST38 epidemiological map was generated using the R software v4.1.1 with different libraries (maptools and ggplot2).

### 2.2. Comparative genomic analyses and in silico characterization of the resistome, plasmidome and virulome

All complete and draft *E. coli* genomes (n=24,102) (last updated in July 2021) were retrieved from GenBank and EMBL databases. The seven MLST alleles that define the ST38 (*adk-4, fumC-26, gyrB-2, icd-25, mdh-5, purA-5*, and *recA-19*) were separately used as queries in BLASTn analyses to recover all currently available ST38 genomes. A total of 376 ST38 genomes recovered in all continents from humans, wild and companion animals, livestock, vegetables, poultry, sewage, fresh, and seawater throughout three decades (1983 – 2020) were used in comparative genomic analyses.

ARG prediction was conducted using the Comprehensive Antibiotic Resistance Database (CARD) (Alcock et al., 2020). The Resistance Gene Identifier (RGI) tool was used for predicting the resistome based on homology and SNP models with the default parameters. The *in silico* detection and plasmid typing based on replicon sequence analysis was performed using the PlasmidFinder web tool (Carattoli et al., 2014). The virulome mining was conducted with ABRicate software package against the *E. coli* virulence database based on VFDB (updated in July 2021) (https://github.com/tseemann/abricate) (Chen et al., 2005). The heatmaps of ST38 resistome, virulome, and plasmidome were drawn using iTOL v4 (Letunic and Bork, 2019).

### 2.3. Phylogenomic analysis

A global-scale phylogenomic reconstruction was performed with the 376 ST38 *E. coli* genomes. These genomes were annotated using Prokka v1.14.6 (Seemann, 2014) and the resultant gff3 files were submitted to Roary v3.13.0 (Page et al., 2015) to determine the core genome. The SNP sites of the concatenated core genes (104,645 bp) were extracted using snp-sites v2.5.1 (Page et al., 2016) and submitted to IQTree v1.6.12 (Minh et al., 2020) to obtain a maximum likelihood SNP tree, which used the model of substitution GTR+F+ASC+R8 and 1000 ultrafast bootstrap replicates. The SNP tree was generated using iTOL v4 (Letunic and Bork, 2019).

## 3. Results and Discussion

### 3.1. Global ExPEC ST38 distribution

A previous study (Manges et al., 2019) established the worldwide distribution of ExPEC lineages based on a systematic revision and meta-analysis considering reports that had performed multilocus sequence typing (MLST) or whole-genome sequencing to genotype *E. coli*. The scenario raised showed that ST38 was in the fourth position on the top 20 ExPEC STs reported in the world. In addition, ST38 occurrence appeared to be lower in North America relative to other continents and absent in Africa (Manges et al., 2019). Here, we analyzed the global occurrence of ExPEC ST38 using the EnteroBase records based on ST38 MLST entries with geographical information (last updated in September, 2021), providing pieces of evidence that enlarge the ST38 prevalence and its worldwide burden.

This meta-analysis revealed that ST38 occurred in countries from all continents, and its prevalence was higher in the USA, Australia, Germany, and the Netherlands, contrasting with the inferences previously established based on reports (Manges et al., 2019). Moreover, this ExPEC lineage occurred in at least nine African countries (Fig. 1).

**Fig. 1.**
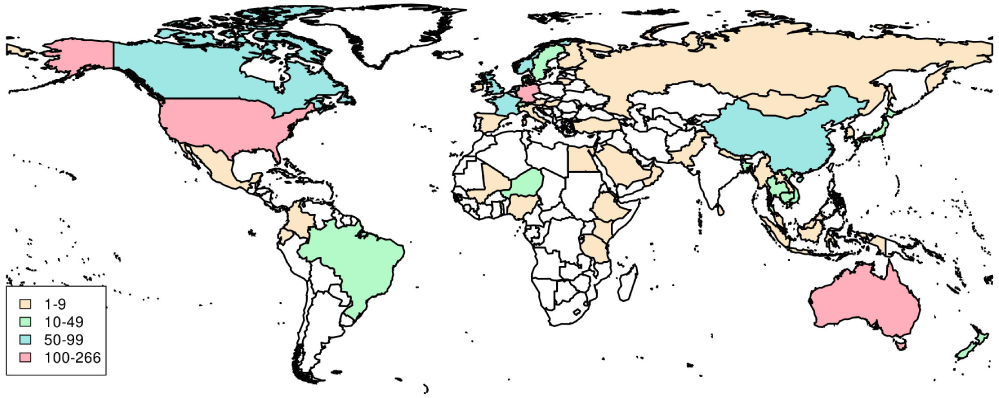
Map of epidemiological distribution of *E. coli* ST38 based on available metadata from EnteroBase (September, 2021).

### 3.2. ST38 Phylogenomics

This study considered 376 ST38 genomes recovered worldwide from humans, animals, and environmental sources persisting for more than three decades (1983-2020). The phylogenomic reconstruction revealed that ST38 genomes grouped in two clusters including genomes from all sources (Fig. 2). Cluster 1 included 32 genomes isolated from 1983 to 2019, while cluster 2 presented several sub-clusters containing the majority of genomes (n=344) that were recovered from 2000 to 2020 (Fig. 2). It was not observed any sub-cluster comprising only genomes from animal and/or environment but there were sub-clusters represented only by genomes from human sources (Fig. 2). Interestingly, despite the overall non-clonal aspect of the ST38, it was observed clonal genomes recovered from humans, animals, and environmental sources in distinct spatiotemporal contexts (Fig. 2). For instance, a Brazilian genome (GCA_019492145), recovered from a urinary tract infection in the Amazon region in 2016 and belonging to a sub-cluster in cluster 1, presented a clonal relationship with genomes from poultry in Australia (2008), animal, and environmental sources recovered in the USA almost 30 years before (1986 and 1993, respectively). In the same way, there was a sub-cluster in cluster 2 comprehending 31 closely related genomes recovered worldwide from distinct sources between 2012 and 2019. The presence of globally distributed ST38 clones points to their epidemiological success in terms of persistence and niche exploration. So far, there were few phylogenomic studies concerning ExPEC lineages. A previous study (Shaik et al., 2017) performed a comparative genomic analysis with the high-risk ExPEC lineages ST131, ST38, ST405, and ST648, in which twelve ST38 genomes recovered from humans were considered. Overall, their study revealed clonality as well as distinctness among strains at the genomic level. Recently, an ST38 genomic analysis revealed the presence of a cluster comprising genomes from humans, wild and companion animals, and animal feed recovered in Asia, Europe, Americas, and Australia between 2013 and 2019, stressing the spreading capacity and ubiquity of some ST38 clones (Lopes et al., 2021). Therefore, these findings altogether demonstrated that ST38 corresponds to a successful zoonotic or zooanthroponosis *E. coli* lineage, probably due to the versatile genome that allows it to persist in animal reservoir hosts and in clinical and natural environments.

**Fig. 2.**
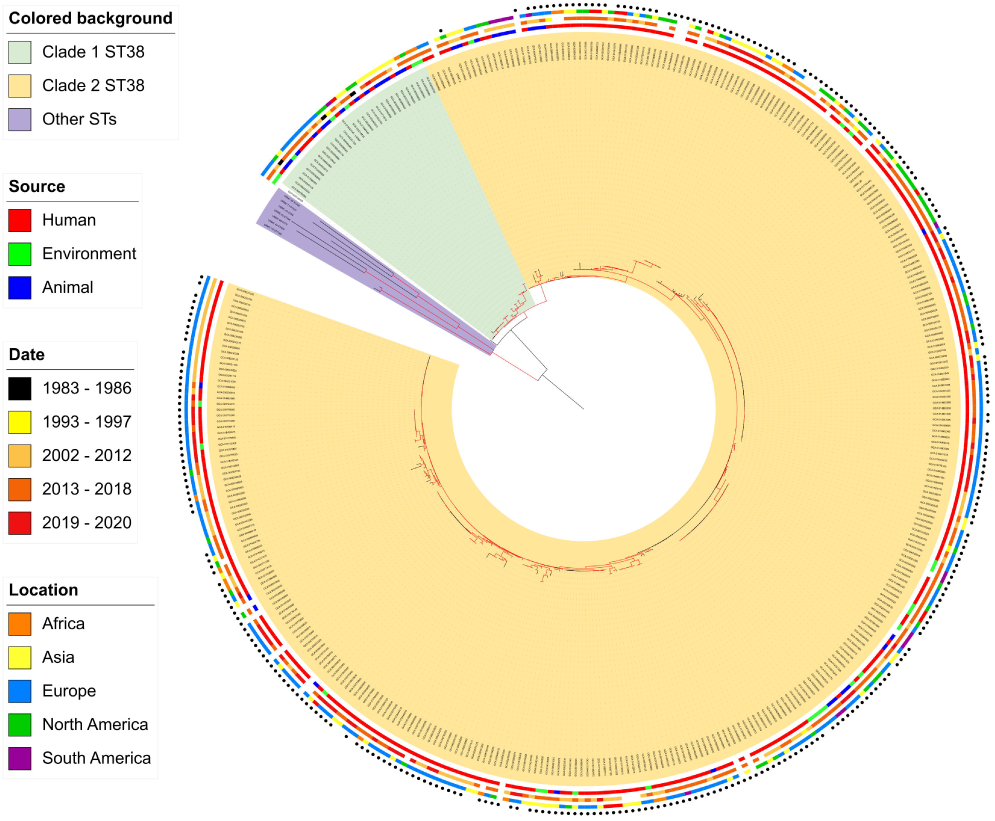
Core SNP tree based on *E. coli* ST38 genomes. Highlighted labels represent the clade 1 (green) and clade 2 (yellow). The circles outside the tree, from innermost to outermost, represent metadata of origin, isolation date, and isolation location, respectively. Genomes featuring the High Pathogenicity Island (HPI) are marked with a black circle. Red branches have bootstrap values greater than 70. An interactive version of the tree is available for specific observations:

### 3.3. ST38 Acquired Resistome

The analysis of the acquired resistome mining revealed the presence of a rich and diverse arsenal of antibiotic resistance genes (ARGs) with a quite heterogeneous distribution among ST38 genomes, independently of their source, genetic relatedness, and spatio-temporal context (Supplementary Fig. S1). Overall, genes involved with resistance to most of the clinically relevant antibiotics for treating *E. coli* infections were found within these genomes, except by those associated with tigecycline and fosfomycin resistance (Magiorakos et al., 2012). A previous study showed a similar acquired resistome profile among other ExPEC lineages such as ST131, ST648, and ST405 (Shaik et al., 2017).

Although the most prevalent ARGs among ST38 were *aph*(*3”)-Ib, aph(6”)-Id, mphA, mphE, qacE*_Δ_*1*, and *sul* (Supplementary Fig. S1), here we focused on ARGs with a remarkable clinical impact such as those coding for ESBLs (*bla*_CTX-M_ and *bla*_CMY-2_), carbapenemases (*bla*_KPC-2_, *bla*_OXA-48_, *bla*_OXA-244_) and metallo-β-lactamases (MBLs) (*bla*_NDM-1_, *bla*_IMP-4_ and *bla*_VIM-4_).

It was found that *bla*_CTX-M_ alleles (*bla*_CTX-M-1, -2, -3, -9, -14, -15, -16, -24, -27, -55_) were dispersed in the great majority of ST38 genomes, corroborating other studies demonstrating the occurrence of *bla*_CTX-M_ in ST38 lineage (Peirano et al., 2012; Pitout, 2012). The *bla*_CTX-M-14_ and *bla*_CTX-M-27_ were the most prevalent among ST38 and were dispersed in several sub-clusters (Supplementary Fig. S1). In fact, *bla*_CTX-M-14_ is one of the most predominant genotype among *Enterobacteriaceae* in the world (Bevan et al., 2017). However, we observed a strong association of *bla*_CTX-M-27_ with a particular subcluster that included ST38 genomes recovered from Vietnam, the USA, France, the Netherlands, Germany, Qatar, Australia, and Brazil (the only genome in this sub-cluster from an animal source). In the case of *bla*_CTX-M-14,_ it was the unique allele found in a sub-cluster comprising genomes recovered between 2006 and 2019 from Asia, Europe, and Oceania from humans and wild animal (one genome from Mongolia). Similarly, *bla*_CTX-M-14_ was also found in a sub-cluster characterized by genomes from the Americas, Asia, Europe and Oceania recovered from all sources between 2012 and 2020. Sporadic genomes (n=4) co-harbored two *bla*_CTX-M_ alleles: *bla*_CTX-M-15_ + *bla*_CTX-M-27_ in a genome from Vietnam; *bla*_CTX-M-15_ + *bla*_CTX-M-9_ from China; *bla*_CTX-M-3_ + *bla*_CTX- M-27_ and *bla*_CTX-M-14_ + *bla*_CTX-M-15_ from France (Supplementary Fig. S1). In general, clonal genomes harbored the same *bla*_CTX-M_ allele. However, eventually, distinct *bla*_CTX- M_ alleles were found among closely related genomes (GCA_016860125 - *bla*_CTX-M -14_; GCA_904863305 - *bla*_CTX-M-15_; GCA_002442325 - *bla*_CTX-M-27_) (Supplementary Fig. S1), corroborating the strong association of *bla*_CTX-M_ with mobile genetic platforms. Moreover, a diversity of *bla*_CTX-M_ alleles, mainly *bla*_CTX-M-14_ and *bla*_CTX-M-15_, was identified among non-human genomes as previously observed elsewhere (Bevan et al., 2017). In comparison with *bla*_CTX-M,_ the ESBL-coding *bla*_CMY-2_ gene was less frequently among ST38, although it was present throughout the clusters (Supplementary Fig. S1).

Carbapenems are drugs of choice to treat infections caused ESBL producing *Enterobacteriaceae* (Gutiérrez- Gutiérrez and Rodríguez-Baño, 2019). Here, genes associated with carbapenem resistance were found among ST38 (*bla*_OXA-48_, *bla*_OXA-244,_ *bla*_KPC-2_, *bla*_NDM-1_, *bla*_IMP-4_, *bla*_VIM-4_), even though in low prevalence, except by the two *bla*_OXA_ genes.

Concerning the carbapenemase-coding genes, the *bla*_OXA-48_ and *bla*_OXA-244_ were found to be the most prevalent and widespread among ST38 (Supplementary Fig. S1). *E. coli* ST38 has been associated with the spread of OXA-48 by clonal expansion (Turton et al., 2016; Pitout et al., 2019). However, our large-scale resistome analysis revealed that *bla*_OXA-48_ distribution in ST38 was not only due to clonal expansion since this gene was found spread among non-clonal genomes from several sub-clusters (Supplementary Fig. S1). Conversely, the *bla*_KPC-2_ gene is rare in ExPEC lineages. Indeed, this gene was found only in two ST38 genomes from Germany (2014) and in a non-clonal genome from Vietnam (2011), all of them from human sources.

In the same way, MBL genes seem to be rare in ST38 since, so far, only *bla*_NDM_ gene had been reported in a few ST38 strains in Asia (Sekizuka et al., 2011; Yamamoto et al., 2011; Shrestha et al., 2017; Huang, et al., 2021). In fact, our resistome mining analysis revealed a sparse occurrence of *bla*_NDM-1_ (n=2; Brazil and Switzerland, 2016), *bla*_IMP-4_ (n=1; China, 1983), and *bla*_VIM-4_ (n=1; France, 2014) among the 376 ST38 genomes. Interestingly, all of these MBL-positive genomes (except for the one from Switzerland) co-harbored a *bla*_CTX-M_ gene: *bla*_CTX-M-15_ + *bla*_NDM-1_ in a Brazilian genome (GCA_019492145); *bla*_CTX-M-14_ + *bla*_CTX-M-15_ + *bla*_VIM-4_ in a genome recovered in France (GCA 009909835); and *bla*_CTX-M-14_ + *bla*_IMP-4_, in an isolate from China (GCA 002223705), all of them associated with human infection cases (Supplementary Fig. S1). Although rare, such association has a relevant impact due to its clinical burden.

### 3.4. ST38 Virulome

The ST38 virulome mining revealed a robust arsenal of virulence-associated genes related to the successful establishment of both intestinal and extraintestinal infections. The ST38 virulome was characterized by genes involved with adherence/aggregation, motility, colonization/invasion, biofilm formation, necroptosis blockage, capsule formation, serum resistance, acid tolerance, iron uptake, intracellular survival, evasion from the host’s immune response, activation of innate immune response and long-term persistence in the urinary tract. However, a heterogeneous distribution of these genes was observed among ST38 regardless of source, phylogenetic relationship, and spatio-temporal context (Supplementary Fig. S2).

Concerning, the virulence genes present in ExPEC pathotypes (UPEC, NMEC, SEPEC, APEC) (Pitout, 2012; Dale and Woodford 2015; Sarowska et al., 2019), ST38 genomes harbored the genes *matF, ibeBC, kpsM, iss, fyuiA, ecpA* and the operons *fim, csg, chu*, and *che*, while other ExPEC virulence genes were not ubiquitous (Supplementary Fig. S2). On the other hand, all ST38 genomes carried genes that play crucial roles in host-pathogen interactions and are associated with adaptability, competitiveness, and colonization capabilities (*aslA, hlyE, motAB, nadAB, malX, artJ, clpV, eaeH, hcp, hofBC, matF, ycfZ, ygdB, b2854/b2972* and the operons *aec, ent, ppd, ecp, flg, flh, fli* and *ycb*) (Peirano et al., 2013; Shaik et al., 2017). These findings reinforce that ST38 is well adapted to survive and proliferate in diverse hosts and environments.

Surprisingly, it was observed a particular and remarkable virulome feature concerning the High Pathogenicity Island (HPI) distribution within ST38 genomes. This island, firstly described in pathogenic *Yersinia* species, encodes the siderophore yersiniabactin (Carniel et al., 1996), and has been considered one of the most relevant virulence factors among ExPEC lineages (Magistro et al., 2017; Shaik et al., 2017; Liu et al., 2018; Galardini et al., 2020, Shan et al., 2021). Besides its pivotal role on iron uptake, the HPI orchestrates various virulence mechanisms and optimizes the overall fitness of ExPEC, such as motility, antibiotic tolerance, and exacerbate inflammatory processes (Magistro et al., 2017; Liu et al., 2018; Galardini et al., 2020, Shan et al., 2021). Considering HPI distribution among ST38, it was observed that this island was absent in cluster 1 and in a specific sub-cluster of cluster 2. Of note is that most of the ST38 genomes in these HPI-missing clusters belonged to animal and environmental sources, contrasting with the strong association and high prevalence of HPI in genomes recovered from humans distributed throughout clade 2 (Fig. 2). Our finding corroborates previous studies on ExPEC lineages, in which HPI was associated with strains causing human infections (Magistro et al., 2017; Shaik et al., 2017; Liu et al., 2018; Galardini et al., 2020, Shan et al., 2021).

### 3.5. ST38 plasmidome

The plasmidome analysis was based on the identification of *rep* genes and their incompatibility group classification (Supplementary Fig. S3). In this way, it was revealed the presence of several plasmids from distinct incompatibility groups among ST38 genomes. The IncFIb group was the most prevalent and evenly distributed in the sub-clusters, and, in fact, it was demonstrated that this plasmid type is spread in several ExPEC lineage such as ST131 (Shaik et al., 2017). In general, there was an abundance of genomes harboring several plasmids (at most 8 per genome). However, two ST38 genomes, one recovered from an animal source (GCA_014712515) and another recovered from human (GCA_015701015), harbored 10 plasmids and nine plasmids, respectively. In addition, some sub-clusters comprised genomes from animal and human sources sharing the same plasmid profile, which included three to five plasmids (Supplementary Fig. S3).

Curiously, the pKPC-CAV1321 plasmid was observed in some ST38 genomes distributed in non-related sub-clusters, recovered from distinct sources and spatio- temporal contexts: one genome from human/Brazil/2016 (GCA_019492145), one from animal/Paraguay/2012 (GCA_016858745), one from wastewater/Czech Republic/2016 (GCA_009790745), and 10 genomes from human/Russia/2012 (Supplementary Fig. S3). The pKPC-CAV1321 had been previously found in *Citrobacter freundii* and *Klebsiella pneumoniae* (Sheppard et al., 2016), and it was recently identified in *Salmonella Choleraesuis* spread in different Brazilian regions (Meneguzzi et al., 2021). However, so far, pKPC-CAV1321 has never been reported in any ExPEC lineage. Interestingly, all pKPC-CAV1321-positive ST38 genomes co-harbored the IncHI2 plasmid. The IncHI2 replicon has been associated with human and animal disease, representing a threat in the context of One Health (Hooton et al., 2021) and, although abundant among *Enterobacteriaceae* (Wong et al., 2016; Matamoros et al., 2017; Zhao et al., 2018), the IncHI2 had never been reported in ST38 and is rare among other ExPEC strains (Shaik et al., 2017).

## 4. Conclusions

Here, we presented the large-scale genomic analyses of the high-risk ExPEC ST38, providing novel insights into its evolutionary mechanisms. It was demonstrated that ST38 is a host-broad pathogen since clones were found efficiently persisting in humans, animals, and natural environments, underlining the global dimension and the zoonotic character of this ExPEC lineage. Probably, the extensive array of resistance genes and the well-equipped arsenal of ExPEC- and IPEC-associated virulence determinants contributed to the establishment of ST38 as a versatile and successful *E*.*coli* lineage.

