## Supplementary Fig. S for "Global genomic epidemiology, resistome, virulome and plasmidome of the Extraintestinal Pathogenic *Escherichia coli* (ExPEC) ST38 lineage"

### SUPPLEMENTARY MATERIAL

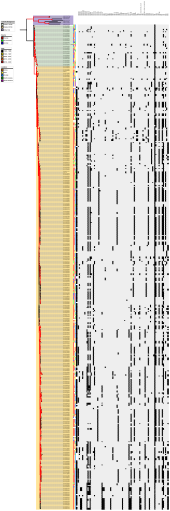

**Supplementary FIG. S1.** Core SNP tree based on E. coli ST38 genomes associated with heatmap of antibiotic resistance genes. Grey and black blocks denote absence and presence of the genes, respectively. Highlighted labels represent the clade 1 (green) and clade 2 (yellow). The colored strips to the right of the tree represent metadata of origin, isolation date, and isolation location, respectively. Genomes featuring the High-Pathogenicity Island (HPI) are marked with a black circle. Red branches have bootstrap values greater than 70.

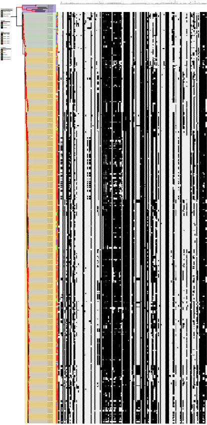

**Supplementary FIG. S2.** Core SNP tree based on E. coli ST38 genomes associated with heatmap of virulence genes. Grey and black blocks denote absence and presence of the genes, respectively. Highlighted labels represent the clade 1 (green) and clade 2 (yellow). The colored strips to the right of the tree represent metadata of origin, isolation date, and isolation location, respectively. Genomes featuring the High-Pathogenicity Island (HPI) are marked with a black circle. Red branches have bootstrap values greater than 70.

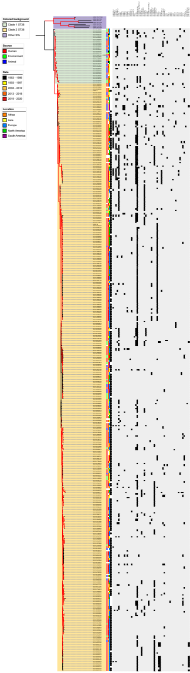


**Supplementary FIG. S3.** Core SNP tree based on E. coli ST38 genomes associated with heatmap of plasmid replicon genes. Grey and black blocks denote absence and presence of the genes, respectively Highlighted labels represent the clade 1 (green) and clade 2 (yellow). The colored strips to the right of the tree represent metadata of origin, isolation date, and isolation location, respectively. Genomes featuring the High-Pathogenicity Island (HPI) are marked with a black circle. Red branches have bootstrap values greater than 70.
